# Comparative Analysis of Structural Alignment Algorithms for Protein-Protein Interfaces in Template-Based Docking Studies

**DOI:** 10.1101/2024.04.03.587755

**Authors:** Fatma Cankara, Nurcan Tuncbag, Attila Gursoy, Ozlem Keskin

## Abstract

Protein-protein interactions are pivotal for various functions within living organisms. Understanding their underlying mechanisms holds significant potential for unraveling cellular processes. There are several methods to identify protein-protein interactions, including but not limited to template-based docking. The power of template docking lies in the template library selection and the quality of structural alignment. Within the scope of our investigation, we specifically delve into the performance of four structural alignment algorithms on one protein interface and four protein structure benchmark sets. This study places particular emphasis on assessing these tools on protein interfaces, composed of non-continuous structure segments, as these interfaces play a crucial role in protein interactions, especially in the context of template-based docking. Notably, our findings indicate that TM-align, despite not being explicitly designed for sequence-order independent alignment, exhibits comparable performance to tools tailored for this purpose while executing in a considerably shorter time frame. Therefore, TM-align emerges as a promising candidate for the crucial structural alignment step in template-docking pipelines.

## 1. Background

Protein-protein interactions (PPIs) perform central roles in the cellular systems of all living organisms [1]. Understanding the interaction mechanism and the characterization of interaction regions of proteins is insightful in revealing the underlying processes driving them [2,3]. Despite the advances in high-throughput interactomics, the gap between structurally resolved protein complexes and known protein interactions is still huge [4]. Protein-protein interaction (PPI) prediction approaches target this challenge through different techniques [5–12], including template-based docking. In template-based docking, a template library of known protein interactions and a target set of proteins, for which their potential interactions are to be predicted, are compared to find structural similarities. Similar targets are then superimposed onto the template to develop a putative protein complex [13]. Shape and physical complementarity are the keys in template-based docking, and the former can be detected via structural alignment [14,15]. Therefore, the performance of template-based approaches is highly dependent on the template library selection and structural alignment quality [16,17]. Some techniques (e.g., Interactome3D [18]) use global structures of the template partners to search for structural similarity with the target, while others (e.g., PRISM [19]) use only the interface region of the template complexes.

In *interface*-template-based docking, the alignment of protein interfaces, i.e. binding sites, emerges as a critical step. Protein structural alignment involves identifying the optimum set of matching points, such as residues or atoms, between two or more proteins based on a score [20–22]. While certain methods incorporate sequence information into the alignment process (e.g., Mican-SQ [23], SPalign [24], TMAlign [25]), many others rely solely on structure-based approaches (e.g., MultiProt [26], SPalign-NS [27], Mican [28], MM-align [29], FTAlign [20–22], US-align [30]). Notably, traditional structure alignment methods are predominantly optimized for aligning global structures rather than specifically targeting protein binding sites or interfaces. In interface-template-based docking, the nature of structural alignment diverges from conventional global alignments to detect fold similarity or identify functional domains. Therefore, interface-template-based docking involves comparing interfaces made of short, non-sequential, non-contiguous patchy segments. In this context, tools (such as iAlign [31], CMAPi [32], and PROSTA-inter [33]), that are specifically developed for this task, are required for the precise alignment of interface regions.

The central component of template-based protein docking is structural alignment, wherein the effectiveness and precision of the alignment process directly influence the quality of generated models [34]. Therefore, we comparatively analyzed the performance of state-of-the-art structural alignment tools in interface (binding site) alignment. Our evaluation focuses on the alignment between interface regions and the global structures of templates intended for use in template-based docking studies. Interfaces are non-contiguous and need to be aligned in a sequence-order independent manner [30,35,36]. While structural alignment tools have been subject to benchmarking in various studies, our approach involves translating the outcomes of these algorithms into the context of template-based protein docking studies by applying them to different scenarios on our interface dataset and carefully selected benchmark sets [37,38].

We evaluated four well-established tools, namely Mican [28], TM-align [25], SPalign-NS [27], and Multiprot [26], chosen based on criteria such as performance, run-time, and availability [22] (Table S1). Our findings indicate that TM-align excels in maximizing the number of matched residues while maintaining acceptable root mean square deviation (RMSD) values. Notably, TM-align demonstrates proficiency in aligning the potential regions rather than trivial random residues. However, we need to note that there are instances where alternative tools outperform TM-align, particularly in cases involving relatively small target structures. Overall, our study provides the trade-offs in the performance of various structural alignment tools and contributes to their applicability in template-based protein-protein docking studies.

## 2. Methods

### 2.1. Datasets

In this study, we used PIFACE, a protein-protein interface dataset by Cukuroğlu et al. [39], and two well-known benchmark sets MALISAM and MALIDUP alongside their sequence-order independent counterparts, MALISAM-ns, and MALIDUP-ns.

The interface dataset, PIFACE, comprises 130,209 interfaces grouped into 22,604 structurally non-redundant clusters. The dataset underwent updates due to certain structures becoming obsolete in the PDB over subsequent years. We used the most crowded clusters as our interface benchmark set (see Supplementary Materials). This dataset has 9,775 interfaces distributed across 78 clusters. Each cluster is represented by a representative structure demonstrating the highest similarity to all its members. We evaluated selected structural alignment tools by examining their ability to align interfaces and overall structures across different scenarios (see 2.2).

Figure 1 illustrates several characteristics of the clusters. On average, clusters have 125.3 member structures, with the most densely populated cluster comprising 193 members and the least densely populated containing 61 members (Figure 1a). The average in-cluster RMSD distribution for selected interfaces is 0.99, as determined by iAlign comparisons (Figure 1b). Moreover, 80% of the proteins within the clusters possess a single PFAM domain, while the predominant secondary structure type identified by CATH classification is alpha-beta (Figure 1c-1d).

**Figure 1.**
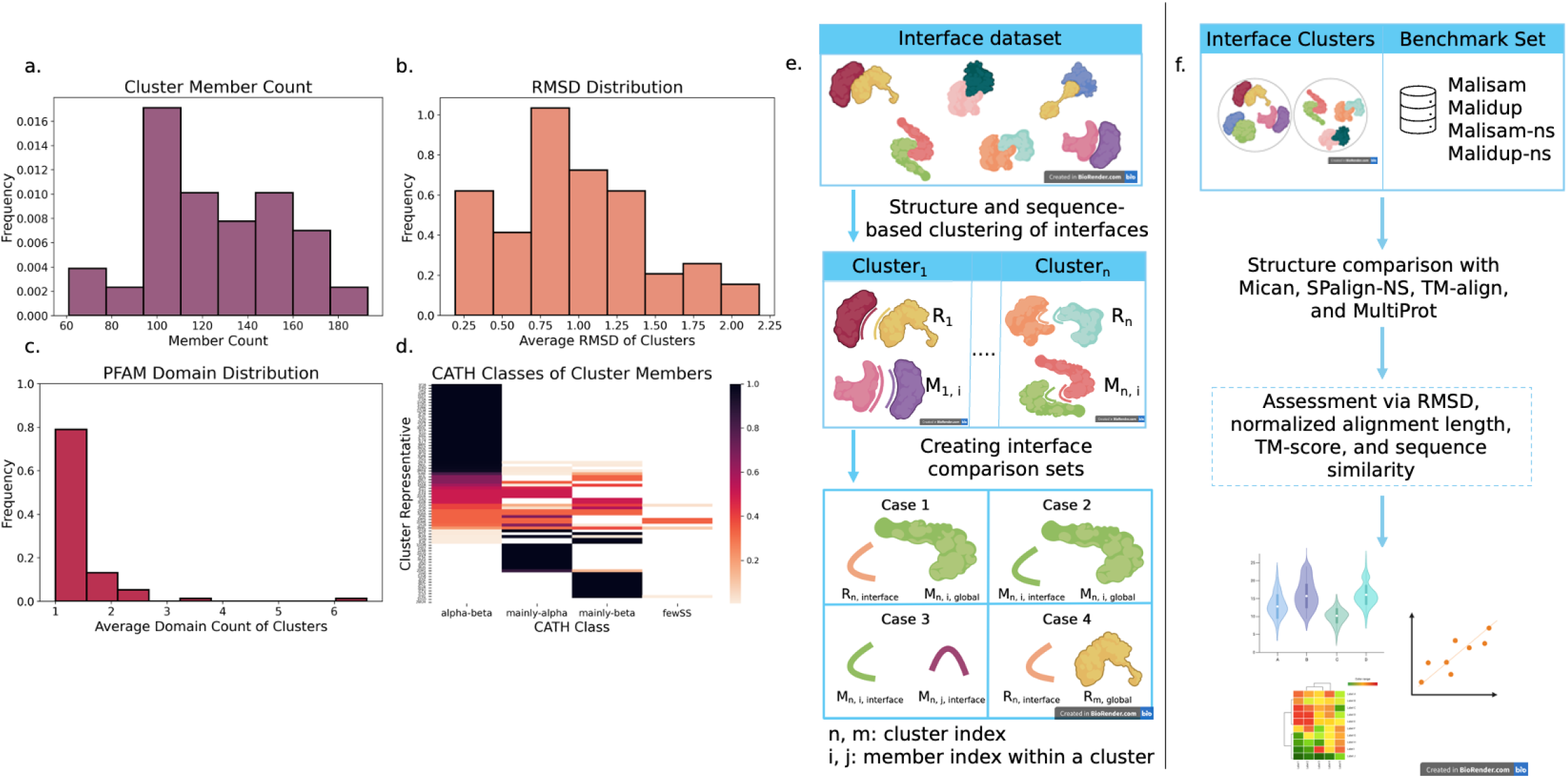
**(a)** Size distribution of 78 structure clusters (max = 193, min = 61, mean = 125.3) **(b)** Average in-cluster RMSD distribution of structure clusters obtained from iAlign comparisons. The average in-cluster RMSD distribution for selected interfaces is 0.99. **(c)** Distribution of average PFAM domain counts of selected structures. On average, selected structures have one PFAM domain. **(d)** CATH classification of selected structures. Darker colors indicate a more homogeneous cluster with similar secondary structures. **(e)** Workflow diagram of the analysis. Stick figures show interfaces; bulk figures show global structures. Index n and m are used for cluster placement, and index *i* and *j* are used for member assignment in the cluster (i.e. M_n,i,global_ is short for ‘the global structure of the *i*th member of the *n*th cluster representative’, while R_n, interface_ is short for ‘the interface structure of the *n*th representative’. Member index is irrelevant for representative structures.). **(f)** Analysis flow for the comparisons. After datasets are processed, structures are aligned based on different scenarios (see 2.2) by implementing four selected structural comparison algorithms. Results are reported by RMSD, normalized alignment length, and TM-score.

On the benchmark side, MALISAM, featuring 130 pairs of structurally analogous motifs, assesses algorithm performance by comparing each structure with its analog [40]. MALIDUP, comprising 241 pairs with non-trivial homology resulting from internal duplication, evaluates selected algorithms through pairwise comparisons [41]. Both sets offer sequence-order independence through MALISAM-ns (130 pairs) and MALIDUP-ns (241 pairs), created by Minami et al. [28] using a multiple-segment permutation technique. Details of these sets can be found in the Supplementary Materials.

### 2.2. Selected Structural Alignment Tools: Multiprot, TMAlign, SPalign-NS, Mican

**MultiProt**, a widely utilized fragment-based tool, employs geometric hashing for aligning multiple proteins and generates partial local solutions. It offers various options, including sequence order dependence or independence, and considers the physicochemical similarity of matched residues.

**TM-align**, a well-established global protein structural alignment tool, utilizes an iterative dynamic programming algorithm for pairwise alignments. Unlike MultiProt, TM-align focuses on global alignments, with TM-score as its objective function.

**SPalign-NS**, an extension of SPalign [24], is a sequence-order independent tool that utilizes the Linear Sum Assignment Problem (LSAP), a combinatorial optimization technique.

**MICAN**, leveraging a geometric hashing technique, detects structurally similar protein pairs while considering secondary structure elements. Notably, MICAN is sequence order independent and addresses over-fragmentation by using a structural similarity score of the Secondary Structure Element (SSE) region instead of individual atoms.

### 2.3. Alignment Evaluation and Quality Metrics

The assessment of alignment quality involves various metrics such as alignment length, RMSD, correctness of aligned regions, and their run-time. We ensure a fair evaluation by normalizing the number of matched residues to the length of the shortest structure.

Our protocols also include eliminating structures with over 80% sequence similarity in global structures and narrowing down the focus to significantly distant pairs, as sequentially similar structures may share topological similarity and may mislead in optimistically reading the alignment results.

## 3. Results

We examined four cases to assess the performance of the structural alignment tools, as illustrated in Figure 1e. Each case delves into a distinct aspect of the interface alignment process, contributing to a comprehensive understanding and interpretation of the results.

***Case 1:*** The structural comparison is between all cluster representatives’ interfaces against the global structures of every member within their respective clusters. This comprehensive comparison serves as a benchmark to assess the efficacy of the tools in identifying interface regions on the global structures of proteins with similar characteristics. In total, 38,900 comparisons are conducted, encompassing the evaluation of 78 cluster representatives and their corresponding member structures.

***Case 2:*** The interface regions of proteins are structurally aligned with their global structures, aiming to gauge their ability to accurately identify their surface region. This analysis encompasses all cluster members, resulting in a total of 19,500 comparisons. Given that the superimposition is onto their structures, we anticipate near-perfect alignments from the alignment tools in this context, this is a positive test case.

***Case 3:*** In the third scenario, we conducted a comprehensive structural comparison by pitting the interfaces of all members against each other within their respective clusters and by comparing every member of a cluster with all the other members in the interface region. Since each cluster is presumed to harbor structurally similar interfaces, this comparison aims to ascertain the ability of the algorithms to align such similar interfaces. The extensive set of comparisons involves a total of 2,557,941 alignments.

***Case 4:*** The interface dataset comprises 78 clusters, each represented by a distinct cluster representative. These representatives are anticipated to exhibit minimal structural similarities to each other due to their distinctiveness, this is a negative control test. To validate this assumption, the interface of each representative structure is aligned with the global structure of all other representatives in the dataset. This analysis is designed to yield trivial alignments, exposing instances of false positives.

We rigorously examine the correctness of alignments through residue-level comparison by comparing matched residues against actual interface residues. In our analysis, structures are aligned on a chain basis, resulting in two alignments for different chains of each dimer structure. One of these alignments is anticipated to represent the correct alignment. The second alignment may yield a high score if the two chains of the target protein have similar interface regions, but this is not guaranteed. Therefore, both alignments are scrutinized to determine the expected alignment pair.

In structural comparisons, we compared aligned residues from global structures to the actual interface residues, determining the ratio of correctly identified residues to all returned residues for Cases 1 and 4. In Case 2, where each structure is compared to itself, no elimination is performed. In Case 3, the chain with the lower RMSD is retained as the correct chain when aligning interfaces. Benchmark comparisons leverage manual alignments from the original studies of MALISAM and MALIDUP. Similar to the interface case, the ratio of correctly identified alignment residues to the alignment residues (reported in the data) is used to assess the correctness of the alignments.

### 3.1. Eliminating less similar chains increases the quality of alignment pairs by correcting aligned residues

In the Methods section (see 2.3), we detailed a process where, for each compared structure, the chain with fewer aligned residues coinciding with interface residues was eliminated, ensuring the selection of the more relevant of the two chains. Figures 2a and 2b depict the impact of this elimination in Case 1 and Case 4 comparisons, respectively. For Case 1, all tools exhibited a distribution skew toward higher ratios after elimination, indicating an improvement in relevance. TM-align showed the most significant increase in correct region matching, followed by Mican and MultiProt. Conversely, in Case 4, where dissimilar structures are compared, a similar pattern was observed, albeit with less difference.

**Figure 2.**
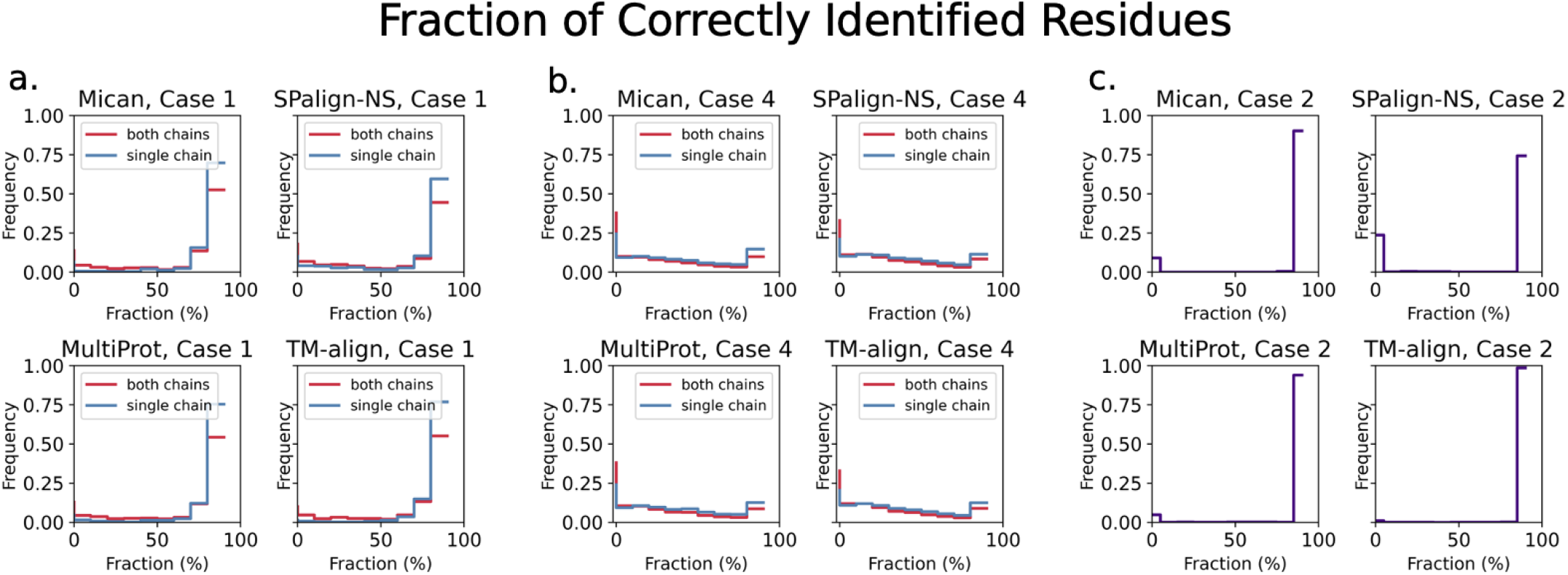
**(a)** Fraction of matched regions for Case 1 comparisons. Red bars indicate the distribution of correctly found interface residue fraction when both of the compared chains are included, while blue bars indicate when the chain with a lower match ratio is filtered out. For all algorithms, the fraction to locate interfaces on global structures increases when the single chain is considered. **(b)** Fraction of matched regions for Case 4 comparisons. **(c)** Fraction of matched regions for Case 2 comparisons. Case 2 involves alignments of interfaces onto their structure; therefore, near-perfect alignments are expected.

In Case 2, where alignments were expected to match interface residues, Figure 2c reveals that a substantial proportion of the comparisons yielded the expected results. SPalign-NS exhibited the highest number of pairs with faulty matching, while TM-align was the most successful.

In benchmark sets, the accuracy of the tools in finding expected alignment residues was assessed by comparing reported residues with manual alignments from the original datasets (Figure S1). MultiProt appeared to report less accurate alignments than the other three tools in both benchmark sets. Mican ranks the best for sequential and non-sequential sets, which particularly stands out for the MALISAM-ns set, while SPalign-NS and TM-align return comparable matches for all cases except for MALIDUP-ns where TM-align shows a similar performance to MultiProt. Mican’s performance for non-sequential sets can be attributed to its design to align non-sequential sets in particular. TM-align’s comparable performance to SP-alignNS, especially when SP-alignNS is used in its sequence-order independent form, indicates that TM-align can achieve alignment results on par with a sequence-independent algorithm.

### 3.2. Comparative analysis of structural alignment methods in interface data indicates TM-align provides better solutions for larger structures

We further refined our analysis by selecting pairs with a sequence similarity of less than 80%. These refined pairs are subsequently utilized as unbiased data in downstream analyses. In Case 1 comparisons, the structural pairs exhibit the highest TM-scores when assessed with TM-align, followed by Mican, SPalign-NS, and MultiProt (refer to Figure 3a). TM-align continues to yield the highest TM-scores in Case 2. In Case 3, SPalign-NS demonstrates the highest TM-score distribution with a mean of 0.70, closely trailed by TM-align and Mican with a marginal difference (mean: 0.68). MultiProt ranks the lowest in Case 3, registering a mean value of 0.53. As anticipated, Case 4 fails to yield meaningful alignments after the filtering steps, where Mican records the highest mean value at 0.32, followed by TM-align, MultiProt, and SPalign-NS, respectively (see Figure 3d). Notably, TM-align consistently provides superior TM-scores for larger structures and interfaces in Case 1 and Case 3 (refer to Tables S1, S2, S3).

**Figure 3.**
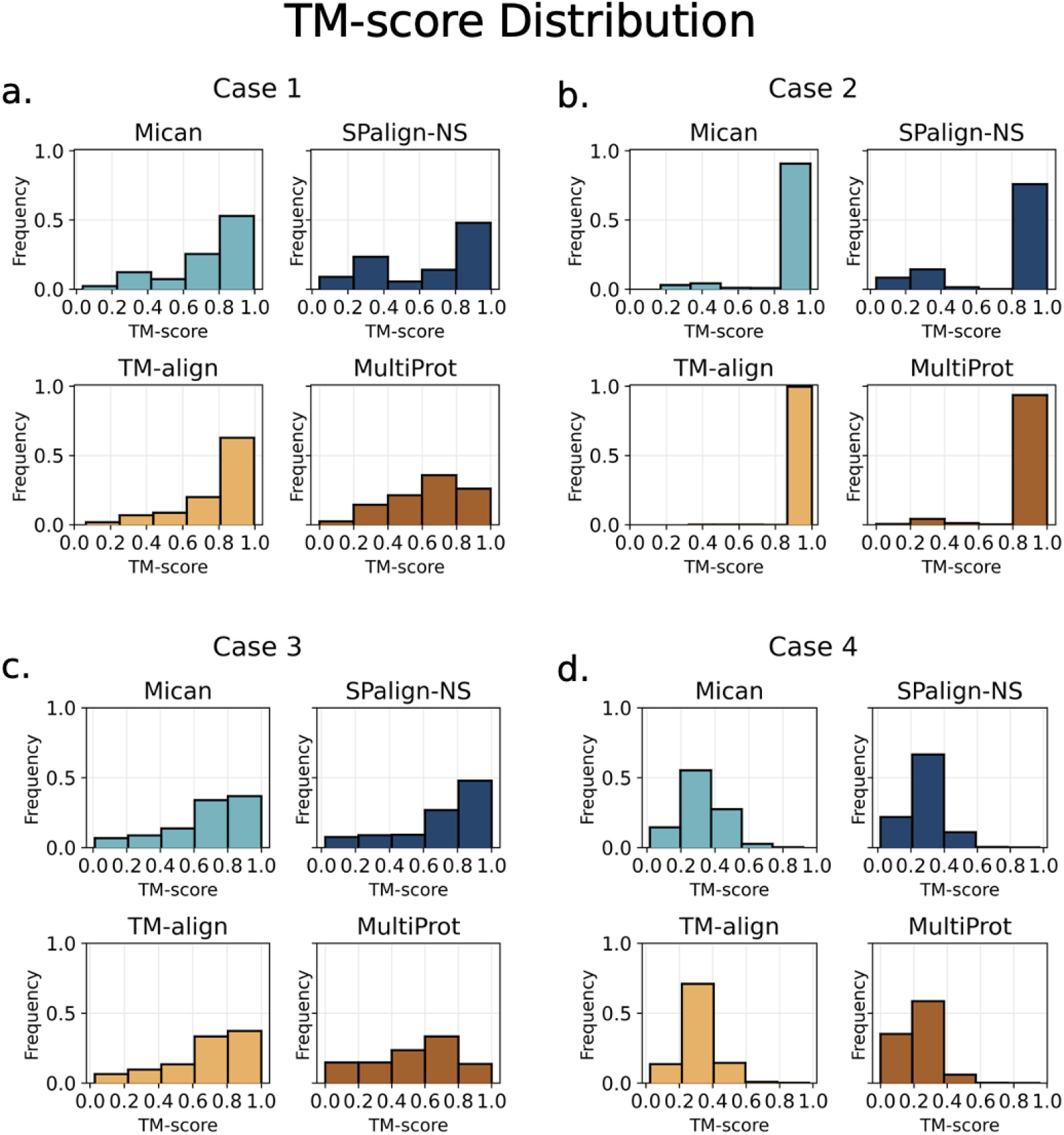
Distribution of TM-scores for **(a)** Case 1 distributions for Mican (mean: 0.73, median: 0.82), SPalign-NS (mean: 0.65, median: 0.78), TM-align (mean: 0.79, median: 0.86) and MultiProt (mean: 0.64, median: 0.70) **(b)** Case 2 distributions for Mican (mean: 0.93 median: 0.98), SPalign-NS (mean: 0.82 median: 1.0), TM-align (mean: 0.99 median: 1.0) and MultiProt (mean: 0.96 median: 1.0) **(c)** Case 3 distributions for Mican (mean: 0.68 median: 0.74), SPalign-NS (mean: 0.7 median: 0.8), TM-align (mean: mean: 0.68 median: 0.75) and MultiProt (mean: 0.53 median: 0.58) and **(d)** Case 4 distributions for Mican (mean: 0.32 median: 0.32), SPalign-NS (mean: 0.28 median: 0.28), TM-align (mean: 0.32 median: 0.31) and MultiProt (mean: 0.23 median: 0.226). Histograms show density distributions.

In the case of benchmark sets, MultiProt exhibits lower performance compared to the other three tools (see Figure S2, Table 1). Mican, SPalign-NS, and TM-align consistently deliver comparable TM-scores, except for MALIDUP-ns and MALISAM-ns cases where Mican performs slightly better. This discrepancy can be attributed to Mican’s fine-tuning for aligning non-sequential structures, highlighting its proficiency in such scenarios compared to other tools. The results obtained from our analysis align with those reported by Wen et al. [22], except for MultiProt, where no data is provided.

**Table 1.**
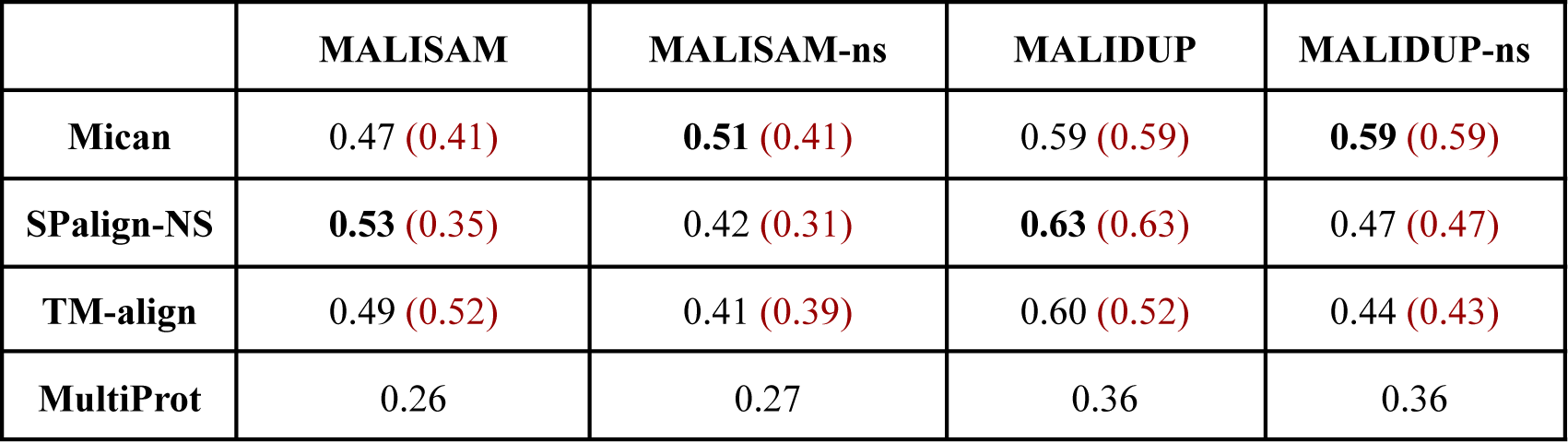
TM-score values for benchmark sets. Results from Wen et al. [22] are indicated in red for Mican, SPalign-NS, and TM-align. MultiProt is not evaluated in Wen et al.

|  | **MALISAM** | **MALISAM-ns** | **MALIDUP** | **MALIDUP-ns** |
| --- | --- | --- | --- | --- |
| **Mican** | 0.47 (0.41) | **0.51** (0.41) | 0.59 (0.59) | **0.59** (0.59) |
| **SPalign-NS** | **0.53** (0.35) | 0.42 (0.31) | **0.63** (0.63) | 0.47 (0.47) |
| **TM-align** | 0.49 (0.52) | 0.41 (0.39) | 0.60 (0.52) | 0.44 (0.43) |
| **MultiProt** | 0.26 | 0.27 | 0.36 | 0.36 |

### 3.3. Analysis of Alignment Accuracy

#### 3.3.1. Evaluation of Interface Set

This section evaluates RMSD and length of the alignments, i.e. the fraction of correctly aligned interface residues, from four protein interface cases and four benchmark sets. Despite Mican’s design for sequence-order independent alignments, it fails to report alignments for a substantial portion of pairs, unlike MultiProt, SPalign-NS, and TM-align. Analyzing the remaining pairs reveals MultiProt’s suboptimal alignment length maximization yet consistently low RMSD values due to its algorithmic constraints. TM-align, especially for sequence-order independent pairs, demonstrates results comparable to Mican and SPalign-NS, exhibiting a lower trade-off between RMSD and normalized alignment length (refer to Figure S3). Furthermore, TM-align proves advantageous in terms of runtime efficiency, delivering similar performance in significantly less time.

In Case 1, the assessment focused on comparing representative interfaces within a cluster, simulating a template-based docking scheme where an interface structure aligns with a template global structure to identify potential interactors. Following filtering steps, which reduced comparisons to 38,900, it was observed that Mican failed to report an alignment in 7.93% of cases. TM-align emerged as the fastest tool, while MultiProt exhibited the lowest RMSD but with a significantly low mean alignment length (Figure 4). It’s essential to highlight that MultiProt employs a heuristic constraint, setting the RMSD threshold to 2 Å for all comparisons in this study. TM-align excelled across multiple metrics, showcasing superior performance in maximizing alignment length, minimizing RMSD, achieving higher TM-scores, and efficiently identifying correct regions in the shortest time. Notably, the sequence similarity among compared structures remained consistent for all tools after the filtering steps, eliminating bias in aligning more similar structures. Average evaluation metrics for Case 1 are detailed in Table S5.

**Figure 4.**
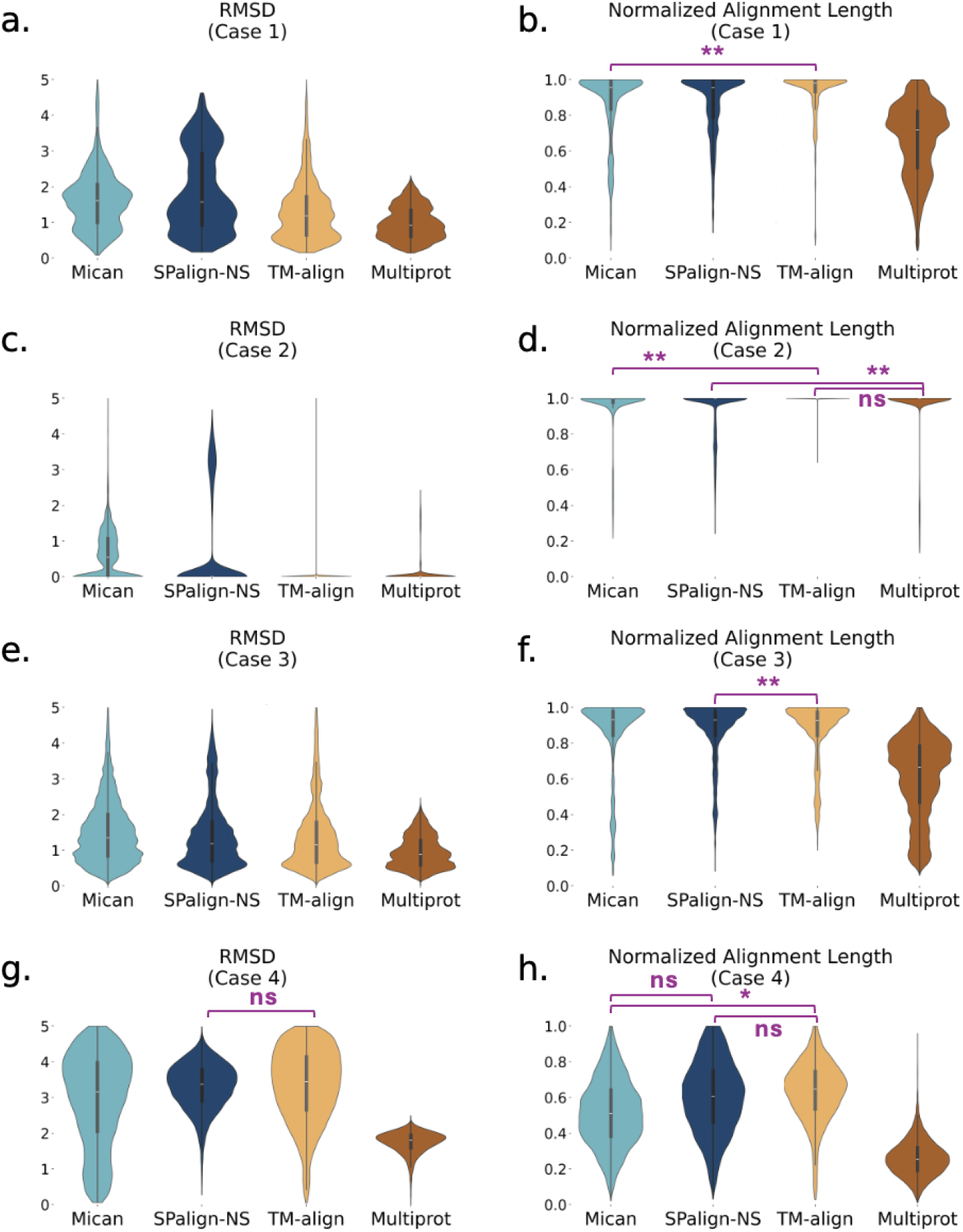
RMSD and normalized alignment length distribution of interface sets for Mican, SPalign-NS, TM-align, and MultiProt. Panels a, c, e, and g show RMSD distribution; while panels a b, d, f, and h show Normalized Alignment Length (Nali) distributions for Case 1, Case2, Case3 and Case 4, respectively. Distributions are given with respective colors as follows: cyan for Mican, navy blue for SPalign-NS, yellow for TM-align, and orange for Multiprot. Lower RMSD with high Nali represents a better alignment. The Kruskal-Wallis Test and Tukey’s HSD Test determine statistical significance, with non-marked pairwise comparisons indicating p-values <= 0.0001, and marked differences having p-values > 0.0001 represented by *** for p-val <= 0.001, ** for p-val <= 0.01, * for p-val <= 0.05, and ns for non-significant cases.

Moving to Case 2, the evaluation focused on comparing member interfaces with their respective global structures. It, therefore, allows the observation of how the selected tools perform in a supposed exact matching case. Unlike Case 1, no filtering was applied to the 19,450 pairs, as structures were compared only to themselves. Mican again failed to return solutions for 7.9% of the pairs (Table S6). TM-align demonstrated superior performance, outperforming other tools across various metrics, even without an RMSD constraint as in Multiprot, and in less time (Figure 4). Notably, SPalign-NS reported the highest RMSD values, while TM-align excelled in accurately identifying correct interface regions.

In Case 3, the assessment involved a comprehensive comparison of interfaces within the same cluster in an all-vs-all manner. MultiProt exhibited the lowest RMSD (0.94 Å) but with a lower mean alignment length (36.63 residues). TM-align showcased superior performance in terms of RMSD and mean alignment length, while Mican aligned the largest number of residues. MultiProt displayed a varying distribution in normalized alignment length, and TM-align stood out with the shortest runtime (Table S7). Excluding abnormal results from MultiProt (RMSD above 100 Å), TM-align continued to demonstrate superior performance.

Case 4 explored the comparison of representative interfaces with the global structure of all other representatives, focusing on identifying false positive alignments. Mican encountered challenges, failing for 11.56% of the companions, while MultiProt faced issues in 0.36% of cases of the total 24,180 comparisons. All tools returned higher RMSD values in this analysis, with a TM-score around 0.3, suggesting trivial alignments. A sequence similarity of approximately 36% indicated minimal structural similarity among the compared structures (Table S8).

#### 3.3.2. Evaluation of the Benchmark Sets

In the subsequent phases of benchmark assessments, a meticulous evaluation of the tools’ performance was conducted on four benchmark sets. In the MALISAM set, the tools successfully provided solutions for all pairs. However, to ensure the fidelity of the analysis, pairs failing to match the anticipated alignment region were excluded. The number of the remaining pairs for each case is reported in Table S9. Notably, MultiProt demonstrated the smallest RMSD, followed by SPalign-NS, TM-align, and Mican (Figure 5). However, when considering the number of aligned residues, MultiProt and Mican fall behind TM-align and SPalign-NS. TM-align and SPalign-NS demonstrated similar TM-score performance, with TM-align excelling in runtime. In this sequence-order dependent set, SPalign-NS and TM-align outperformed Mican and MultiProt.

**Figure 5.**
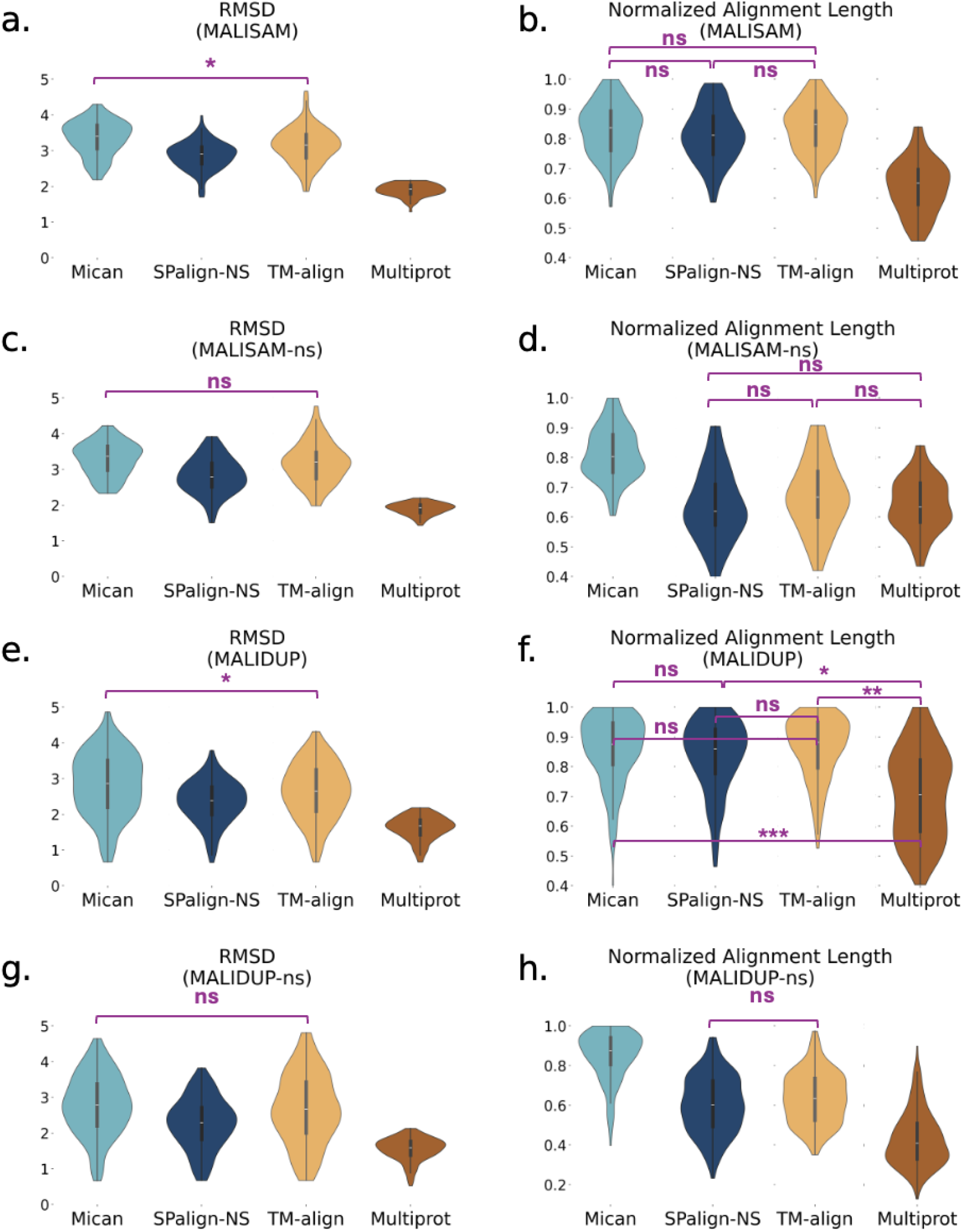
RMSD and normalized alignment length distribution of benchmark sets for Mican, SPalign-NS, TM-align, and MultiProt. Panels a, c, e, and g show RMSD distribution; while panela b, d, f, and h show Normalized Alignment Length (Nali) distributions for Malisam, Malisam-ns, Malidup, and Malidup-ns respectively. Distributions are given with respective colors as follows: cyan for Mican, navy blue for SPalign-NS, yellow for TM-align, and orange for Multiprot. Lower RMSD with high Nali represents a better alignment. The Kruskal-Wallis Test and Tukey’s HSD Test determine statistical significance, with non-marked pairwise comparisons indicating p-values <= 0.0001, and marked differences having p-values > 0.0001 represented by *** for p-val <= 0.001, ** for p-val <= 0.01, * for p-val <= 0.05, and ns for non-significant cases.

In the MALISAM-ns set, only comparisons with correctly aligned regions were considered when calculating average values, similar to the approach taken in the MALISAM case (Table S10). Mican showed better performance than other tools, except in RMSD. TM-align aligned a comparable number of residues but displayed a shortfall in normalized alignment length. This observation implies that TM-align tends to yield consistent RMSD values for aligning larger structures, given the normalization process. TM-align’s ability to perform similarly to tools designed for sequence-order independence indicates its adaptability. Mican outperformed in finding correct alignment regions, while SPalign-NS and TM-align performed similarly. Overall, SPalign-NS and TM-align showed similar performance, except for lower RMSD in SPalign-NS. Mican excelled in normalized alignment length, TM-score, and finding correct alignment regions. Mican, SPalign-NS, and TM-align show comparable RMSD values with a similar alignment length.

The MALIDUP set revealed a noticeable alignment of larger fragments compared to the MALISAM set across all four selected tools, with TM-align leading in this context. TM-align excelled in aligning larger fragments, as evidenced by superior RMSD and TM-score (Figure 5). In contrast, MultiProt, while presenting lower RMSD values, failed to align long regions compared to others. Remarkably, TM-align achieved this commendable performance with the shortest alignment time of 0.1 s per alignment (Table S11).

Concluding with the MALIDUP-ns set, another sequence-order independent set derived from MALIDUP, Mican, and SPalign-NS, tuned for sequence-order independent alignment, showed comparable performance. TM-align also exhibited similar performance in mean RMSD and mean alignment length. Mican outperformed them in TM-score and the fraction of correctly aligned regions, while SPalign-NS and TM-align were similar in these metrics (Table S12, Figure 5).

### 3.4. TM-align and MultiProt comparison reveals TM-align aligns longer structures with comparable RMSD

PRISM [5] employs a template-based docking approach to identify protein interactors by assessing interaction region similarities, utilizing MultiProt as its chosen structural alignment algorithm. In order to compare the performance of MultiProt and TM-align, we evaluated a subset of 8574 pairs for which both tools give results. In Case 1 comparisons, TM-align demonstrated the ability to align identical structures over longer regions with comparable RMSD and a higher correct match ratio (Figure 6). Specifically, the average RMSD values for Case 1 were 1.29 for TM-align and 0.96 for MultiProt. Despite similar RMSD values, TM-align exhibited a significantly higher normalized alignment length (68.77 vs. 49.25 for MultiProt). The matching accuracy remained similar, with values of 89.85 for TM-align and 88.33 for MultiProt, suggesting that MultiProt aligns correct regions but at a shorter length. Benchmark comparisons, detailed in Figure S4, consistently demonstrated that TM-align aligns longer regions with either comparable or superior accuracy in identifying correct matches.

**Figure 6.**
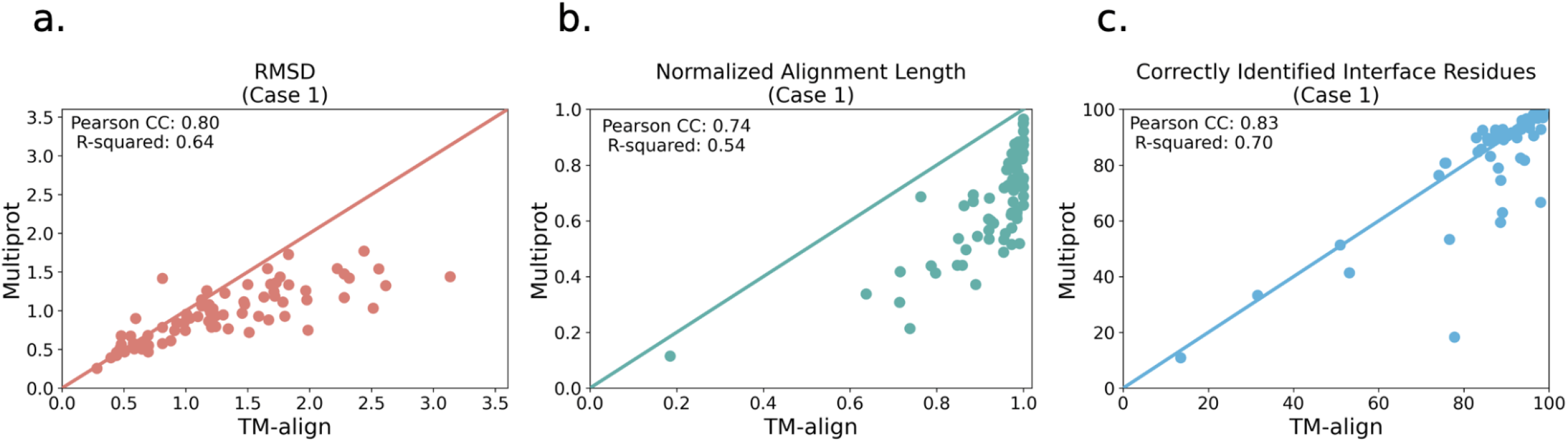
RMSD, normalized alignment length, and the fraction of correctly aligned residue comparisons for the subset pairs of interface sets for which both TM-align and MultiProt give results.

## 4. Conclusions

Structural alignment is a critical step in template-based PPI docking, as the efficacy and accuracy of the alignment influence the quality of the models. Consequently, the present investigation undertook a comprehensive comparison of diverse structural alignment tools to evaluate their proficiency in assessing binding site alignment. We considered different scenarios, including the mimicking of a template-based docking scheme wherein template interfaces were aligned to established targets (Case 1). In general, TM-align appeared to perform similarly to non-sequential alignment tools Mican and SPalign-NS, even though its algorithm does not necessarily consider sequentiality. Notably, TM-align consistently yielded extended alignments accompanied by relatively lower RMSD values, indicative of the non-random matching of aligned patches. The comparable performance of TM-align extended to non-sequential datasets such as MALISAM-ns and MALIDUP-ns, underscoring its suitability in diverse alignment contexts within a template-based docking scheme. Due to its restricted RMSD threshold of 2Å, MultiProt posed a hindrance to direct comparison with the other three tools. Nevertheless, a comprehensive assessment considering normalized alignment length, time, and TM-score exposed TM-align’s superiority across these metrics. In summation, TM-align consistently exhibited longer alignments with lower RMSD values and higher TM-scores within a significantly abbreviated timeframe. Consequently, it emerges as a discerning choice for structural alignment in template-based docking studies.

## Acknowledgment

This project has been partially funded by TUBITAK 120C120 project. Fatma Cankara is supported by the TUBITAK BIDEB 2211/A National PhD Scholarship Program.

## Supporting Information

Supporting Information on the selected algorithms, datasets and additional figures are provided in Supplementary Files. Supplementary Table S1: Summary about the selected tools; Supplementary Table S2: The mean length of global and interface structures in the pairs with different TM-score thresholds for Case 1; Supplementary Table S3: The mean length of global and interface structures in the pairs with different TM-score thresholds for Case 2; Supplementary Table S4: The mean length of global and interface structures in the pairs with different TM-score thresholds for Case 3; SupplementaryTable S5: Comparative metrics for Case 1 comparisons; SupplementaryTable S6: Comparative metrics for Case 2 comparisons; Supplementary Table S7: Comparative metrics for Case 3 comparisons; Supplementary Table S8: Comparative metrics for Case 4 comparisons; Supplementary Table S9: Comparative metrics for MALISAM comparisons; Supplementary Table S10: Comparative metrics for MALISAM-ns comparisons; Supplementary Table S11: Comparative metrics for MALIDUP comparisons; Supplementary Table S12: Comparative metrics for MALIDUP-ns comparisons. Supplementary Figure S1: The distribution of the number of correctly matching residues for benchmark sets; Supplementary Figure S2: Distribution of TM-scores for benchmark sets; Supplementary Figure S3: Scatter plot for normalized alignment length vs. RMSD for the interface and benchmark set analysis; Supplementary Figure S4: RMSD, normalized alignment length, and the fraction of correctly aligned residue comparisons for the subset pairs of benchmark sets for which results are given by both TM-align and MultiProt.

## Author Contributions

F.C., N.T., A.G., and O.K. designed and conceptualized the project. F.C. analyzed data, and prepared tables, and figures. F.C., N.T., A.G., and O.K. wrote and edited the manuscript. All the authors reviewed and approved the final manuscript.

