## Supplementary Materials for "Comparative Analysis of Structural Alignment Algorithms for Protein-Protein Interfaces in Template-Based Docking Studies"

#### 1. **Background**

**Table S1. Summary of the selected tools.**

|  | **Available link** | **Rigid superposition** | **Represent** | **Search algorithm** | **Pairwise/**  **Multiple** | **Optimization based on** | **Year** |
| --- | --- | --- | --- | --- | --- | --- | --- |
| **Multiprot** | http://bioinfo3d.cs.tau.ac.il/MultiProt/php.php | Yes | C-alpha | Geometric hashing | Multiple | RMSD values and alignment length | 2004 |
| **TM-align** | https://zhanggroup.org/TM-align/ | Yes | C-alpha | Dynamic programming | Pairwise | TM-score | 2005 |
| **MICAN** | http://landscape.tbp.cse.nagoya-u.ac.jp/MICAN/ | Yes | SSE | Geometric hashing | Pairwise | sTMscore | 2013 |
| **SP-alignNS** | https://sparks-lab.org/downloads/ | Yes | C-alpha | Dynamic programming | Pairwise | SP-score | 2015 |

#### **2.1. Datasets**

Interface cluster representatives are given below:

1EBD_AB, 1F28_AB, 1FB1_AE, 1FRS_AC, 1FSK_FI, 1GQM_KL, 1H28_AD, 1HA7_QR, 1IAO_AB, 1IR2_7Y, 1JXN_CD, 1LWF_AB, 1MPS_HM, 1O1D_1Z, 1RCX_OT, 1RD8_AE, 1SQ7_AB, 1SWJ_AB, 1T0A_AC, 1UQR_EK, 1UV6_GH, 1UZD_IK, 1VMK_AC, 1ZBB_GH, 1ZUM_AC, 2AUQ_AB, 2BE7_CF, 2CE4_AB, 2F36_CD, 2FG8_CD, 2FKZ_CD, 2GMR_HL, 2NVL_CE, 2O5I_MN, 2OIZ_BD, 2OM1_dh, 2P2C_KL, 2PMI_CD, 2PPY_BF, 2QUR_AB, 2VDI_CE, 2WUH_BD, 2X0N_PR, 2XOK_CD, 2Y81_AB, 2ZL4_GN, 3AZG_CE, 3AZI_BG, 3BXC_CD, 3DOC_CD, 3DPY_AB, 3FCP_DG, 3FDU_AC, 3G0B_AB, 3GVI_AD, 3HFA_UV, 3KX9_PU, 3L7Q_GI, 3L9E_AB, 3LC7_QR, 3MFE_1U, 3MH4_AB, 3N5Z_AB, 3NWL_AB, 3OAA_Zc, 3OID_BD, 3P7X_BD, 3PCC_DP, 3PCE_MN, 3QPB_EF, 3R45_AB, 3SVR_AC, 3TA1_EF, 3TPU_AB, 3TV3_HL, 3UBE_GJ, 3UBE_HL, 4A3L_CK

Members for each cluster can be accessed at the following link: <https://interactome.ku.edu.tr/piface/depositor/finalInterfaceClusters_2013_January_24_.txt>

In addition to the interface sets, two commonly used benchmark sets for structural comparison are used to assess the performance of selected tools. Two of these sets, MALISAM and MALIDUP, are manually created sequence-order dependent datasets, while the other two, MALISAM-ns and MALIDUP-ns, are sequence-order independent. MALISAM (manual alignments for structurally analogous motifs) consists of 130 pairs of structural analogs. Analogy refers to the similarity between protein structures that are not a result of evolutionary relationships; rather, it results from converging to similar structures due to a limited number of energetically favorable ways to pack secondary structural elements [[1]](https://paperpile.com/c/xK91md/AZR7). Each structure in this set is compared against the given analog structure to assess the performance of selected algorithms. In total, 130 pairwise comparisons are performed for the MALISAM set.

MALIDUP (manual alignments of duplicated domains) contains 241 pairs created by structural alignments with non-trivial homology [[2]](https://paperpile.com/c/xK91md/K4eHz). It comprises homologous domains originating from internal duplication within the same polypeptide chain. The duplication process eventually produces sequentially and functionally different proteins. Structures in these sets are given as pairs. Thus they are compared with the corresponding protein.

The sequence-order independent pairs are created from these two sequence-order dependent sets using multiple segment permutation technique artificially by Minami et al. [[3]](https://paperpile.com/c/xK91md/GCt2q). MALISAM-ns is created from the MALISAM dataset and thus contains 130 pairs. Similarly, MALIDUP-ns is created from the MALIDUP set and contains 241 pairs.

#### **3.1. Eliminating less similar chains increases the quality of alignment pairs by correcting aligned residues**

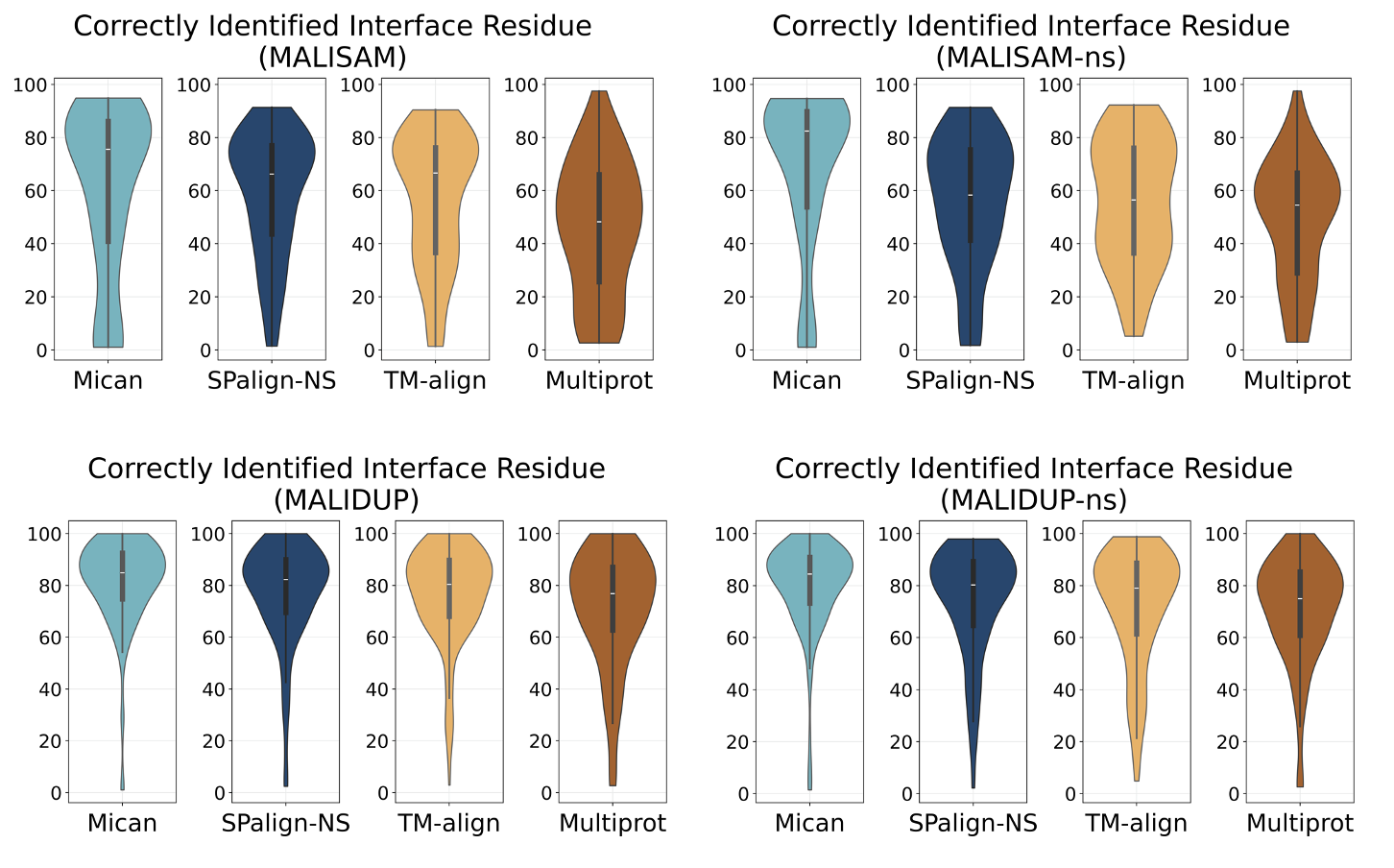

**Figure S1.** The distribution of the number of correctly matching residues for benchmark sets. **(a)** MALISAM distributions for Mican (mean: 60.19 median: 75.57), SPalign-NS (mean: 58.98 median: 66.26), TM-align (mean: 56.72 median: 66.67), and MultiProt (mean: 46.26 median: 48.28) **(b)** MALISAM-ns distributions for Mican (mean: 68.44 median: 82.42), SPalign-NS (mean: 55.86 median: 58.33), TM-align (mean: 55.5 median: 56.52) and MultiProt (49.38 median: 54.55) **(c)** MALIDUP distributions for Mican (mean: 81.48 median: 84.9), SPalign-NS (mean: 77.22 median: 82.23), TM-align (mean: 76.55 median: 80.0) and MultiProt (mean: 70.16 median: 76.67) **(d)** MALIDUP-ns distributions for Mican (79.53 median: 84.44), SPalign-NS (mean: 74.3 median: 80.26), TM-align (70.85 median: 78.39) and MultiProt (mean: 70.69 median: 75.0). Numbers are reported by eliminating the pairs that did not return common residues with the manual alignments for the alignment tool of interest.

#### **3.2. Comparative analysis of structural alignment methods in four cases indicates TM-align provides better solutions for larger structures**

**Table S2.** The mean length of global and interface structures in the pairs with different TM-score thresholds for Case 1.

|  | **Mican < 0.5** | | **MultiProt < 0.5** | | **SPalign-NS < 0.5** |
| --- | --- | --- | --- | --- | --- |
| **TM-align > 0.5** | 452 pairs | | 1326 pairs | | 0 pairs |
|  | Interface | Global | Interface | Global |  |
|  | **41.87** | **259.65** | **84.89** | **264.01** |  |
|  | **Mican > 0.5** | | **MultiProt > 0.5** | | **SPalign-NS > 0.5** |
| **TM-align < 0.5** | 107 pairs | | 84 pairs | | 0 pairs |
|  | Interface | Global | Interface | Global |  |
|  | 41.27 | 202.87 | 49 | 219.78 |  |

**Table S3.** The mean length of global and interface structures in the pairs with different TM-score thresholds for Case 2.

|  | **Mican < 0.5** | | **MultiProt < 0.5** | | **SPalign-NS < 0.5** | |
| --- | --- | --- | --- | --- | --- | --- |
| **TM-align > 0.5** | 1298 pairs | | 1125 pairs | | 4572 | |
|  | Interface | Global | Interface | Global | Interface | Global |
|  | 35.54 | **273.112** | 37.73 | **303.20** | 44.7 | **297** |
|  | **Mican > 0.5** | | **MultiProt > 0.5** | | **SPalign-NS > 0.5** | |
| **TM-align < 0.5** | 18 pairs | | 15 pairs | | 7 | |
|  | Interface | Global | Interface | Global | Interface | Global |
|  | **55.56** | 254.39 | **56.67** | 262.4 | **57.71** | 252.42 |

**Table S4.** The mean length of global and interface structures in the pairs with different TM-score thresholds for Case 3.

|  | **Mican < 0.5** | | **MultiProt < 0.5** | | **SPalign-NS < 0.5** |
| --- | --- | --- | --- | --- | --- |
| **TM-align > 0.5** | 17252 pairs | | 113183 pairs | | 0 pairs |
|  | Interface1 | Interface2 | Interface1 | Interface2 |  |
|  | **80.98** | **75.92** | **84.16** | **84.52** |  |
|  | **Mican > 0.5** | | **MultiProt > 0.5** | | **SPalign-NS > 0.5** |
| **TM-align < 0.5** | 12632 pairs | | 6514 pairs | | 0 pairs |
|  | Interface1 | Interface2 | Interface1 | Interface2 |  |
|  | 38.14 | 39.01 | 59.47 | 46.82 |  |

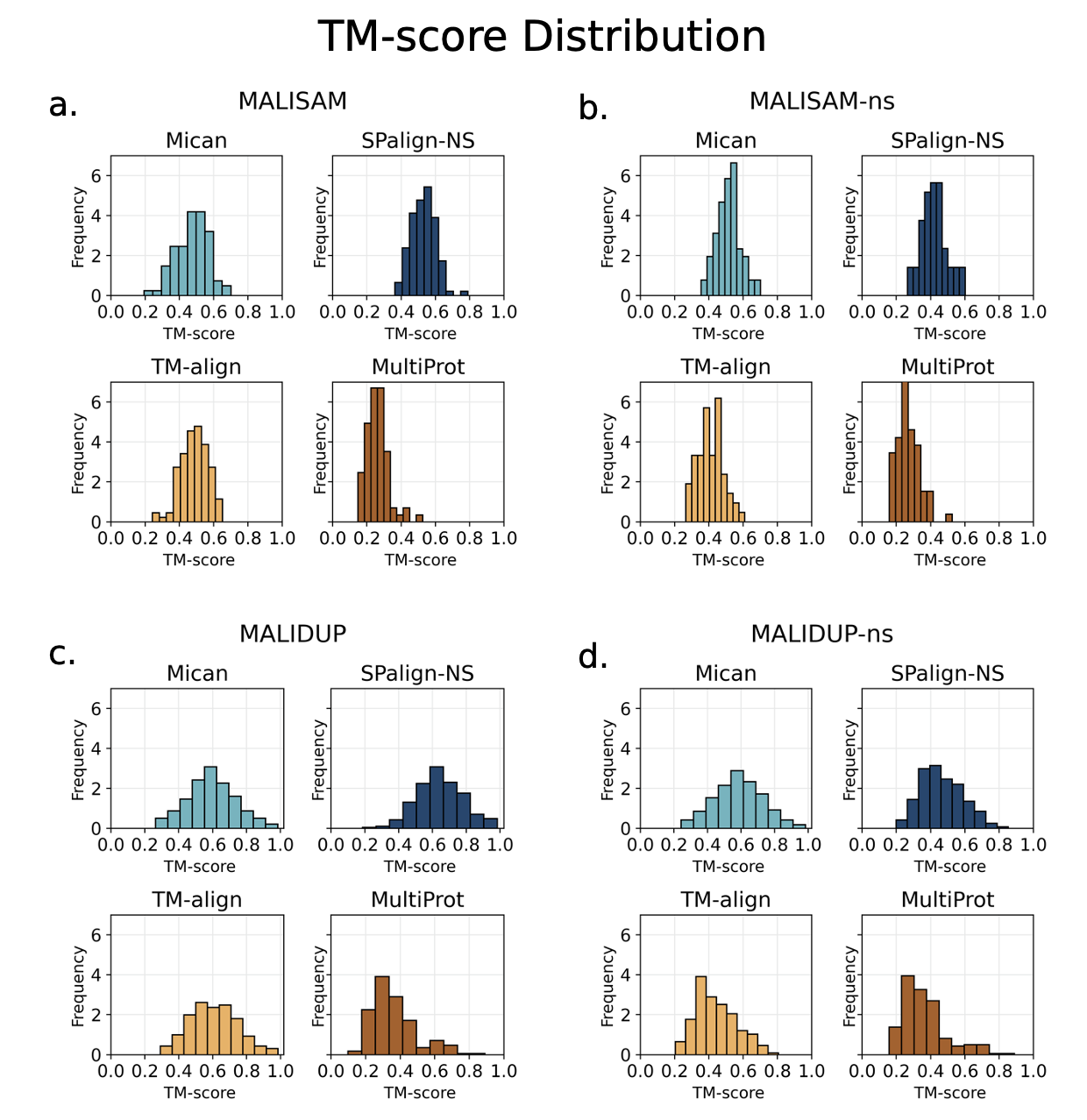

**Figure S2:** Distribution of TM-scores for **(a)** MALISAM distributions for Mican (0.47 median: 0.48), SPalign-NS (mean: 0.53 median: 0.53), TM-align (mean: 0.49 median: 0.49) and MultiProt (mean: 0.26 median: 0.26) **(b)** MALISAM-ns distributions for Mican (mean: 0.51 median: 0.51), SPalign-NS (mean: 0.42 median: 0.42), TM-align (mean: 0.41 median: 0.41) and MultiProt (mean: 0.27 median: 0.26) **(c)** MALIDUP distributions for Mican (mean: 0.59 median: 0.6), SPalign-NS (mean: 0.63 median: 0.64), TM-align (mean: 0.60 median: 0.6) and MultiProt (mean: 0.36 median: 0.33) **(d)** MALIDUP-ns distributions for Mican (mean: 0.59 median: 0.59), SPalign-NS (mean: 0.47 median: 0.46), TM-align (0.44 median: 0.42) and MultiProt (0.36 median: 0.33)

#### **3.3. Analysis of Alignment Accuracy**

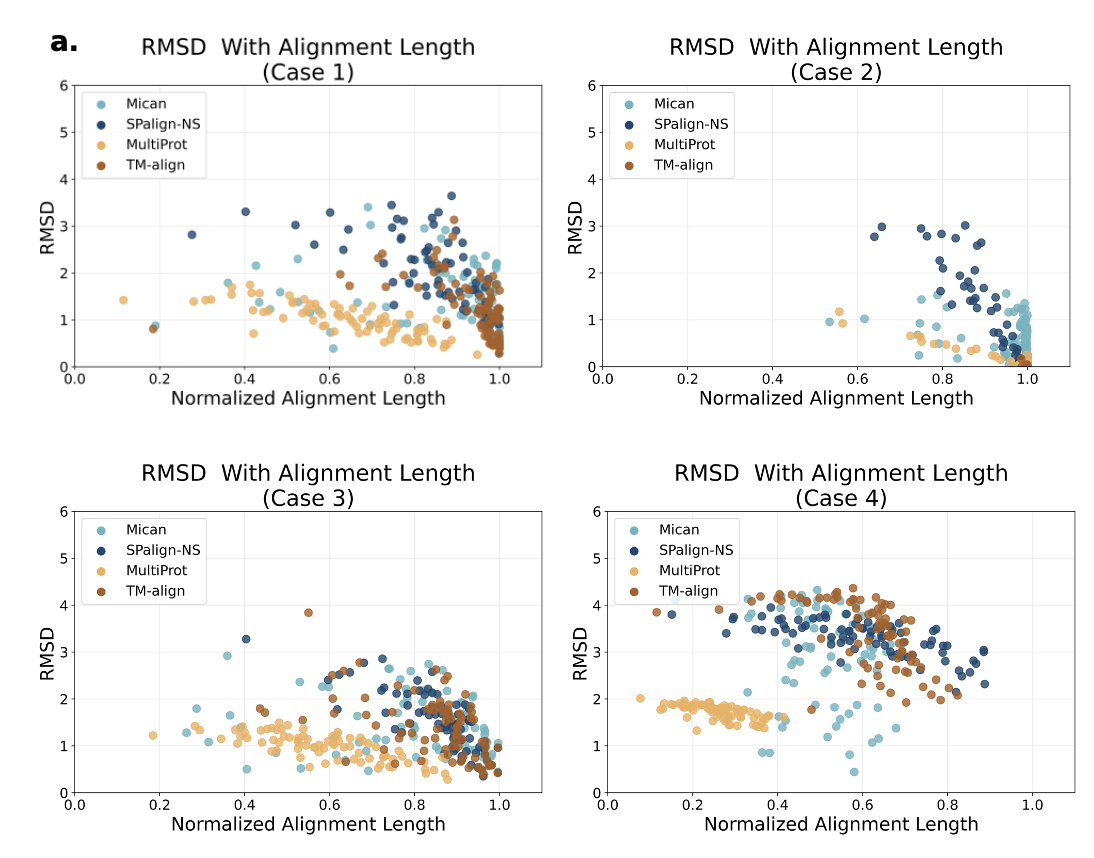

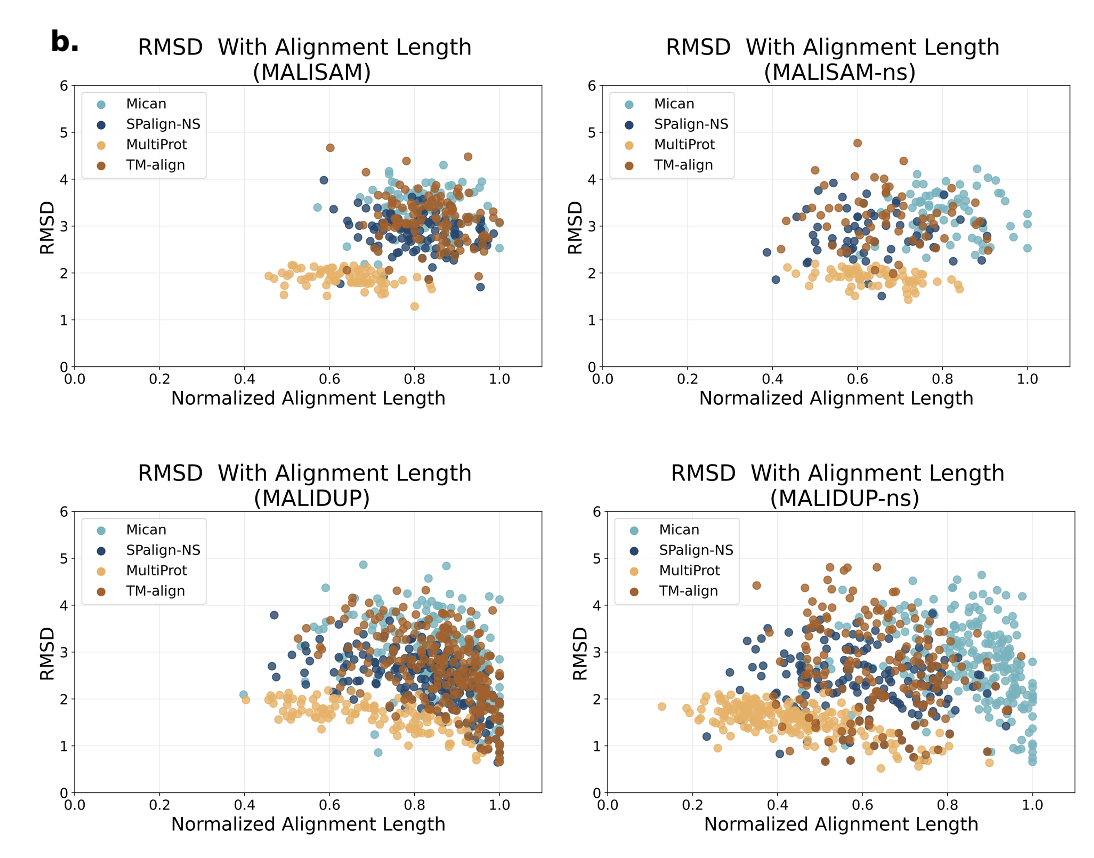

**Figure S3:** Scatter plot for normalized alignment length vs. RMSD for the **(a)** interface and **(b)** benchmark set analysis. Alignment lengths are normalized by the shortest of the compared structures, which is either the size of the interface when compared to a global structure or the shorter of interfaces when two interfaces are compared.

**Table S5:** Comparative metrics for Case 1 comparisons

| **Case 1** | **Mican** | **SPalign-NS** | **TM-align** | **MultiProt** |
| --- | --- | --- | --- | --- |
| **Number of aligned pairs** | 11,025 | 11,500 | 11,188 | 11,355 |
| **RMSD** | 1.62 | 1.87 | 1.31 | **0.98** |
| **Alignment length** | 63.39 | 60.37 | **65.49** | 46.06 |
| **Normalized alignment length (%)** | 86.6 | 86.6 | **93.1** | 66.3 |
| **TM-score** | 0.73 | 0.65 | **0.79** | 0.64 |
| **Fraction of correctly aligned regions** | 81.71 | 73.11 | **88.48** | 85.90 |
| **Sequence similarity between structures** | 0.51 | 0.51 | **0.52** | **0.52** |
| **Fraction of comparisons for which a solution is not given for (%)** | 7.93 | **0** | **0** | **0** |
| **Time per alignment (s)** | 0.17 | 0.26 | **0.08** | 0.65 |

**Table S6.** Comparative metrics for Case 2 comparisons

| **Case 2** | **Mican** | **SPalign-NS** | **TM-align** | **Multiprot** |
| --- | --- | --- | --- | --- |
| **Number of aligned pairs** | 17,905 | 19,450 | 19,450 | 19,450 |
| **RMSD** | 0.63 | 0.77 | **0.01** | 0.02 |
| **Alignment length** | **71.33** | 66.33 | 69.38 | 68.06 |
| **Normalized alignment length (%)** | 95.04 | 93.58 | **99.90** | 96.11 |
| **TM-score** | 0.93 | 0.82 | **1.0** | 0.96 |
| **Fraction of correctly aligned regions** | 89.13 | 74.11 | **98.78** | 94.53 |
| **Sequence similarity between structures** | **0.51** | **0.51** | **0.51** | **0.51** |
| **Fraction of comparisons for which a solution is not given for (%)** | 7.9 | **0** | **0** | **0** |
| **Time per alignment (s)** | 0.31 | **0.29** | 0.31 | 0.85 |

**Table S7.** Comparative metrics for Case 3 comparisons.

| **Case 3** | **Mican** | **SPalign-NS** | **TM-align** | **MultiProt** |
| --- | --- | --- | --- | --- |
| **Number of aligned pairs** | 794,979 | 757,818 | 759,693 | 760,232 |
| **RMSD** | 1.53 | 1.40 | 1.41 | **0.94** |
| **Alignment length** | **57.92** | 56.49 | 56.77 | 39.63 |
| **Normalized alignment length (%)** | 0.84 | **0.87** | 0.86 | 0.61 |
| **TM-score** | 0.68 | **0.70** | 0.68 | 0.53 |
| **Sequence similarity between structures** | **0.51** | **0.51** | **0.51** | **0.51** |
| **Fraction of comparisons for which a solution is not given for (%)** | 9.9 | **0** | **0** | 0.02 |
| **Time per alignment (s)** | 0.064 | 0.071 | **0.022** | 0.598 |

**Table S8.** Comparative metrics for Case 4 comparisons

| **Case 4** | **Mican** | **SPalign-NS** | **TM-align** | **Multiprot** |
| --- | --- | --- | --- | --- |
| **Number of aligned pairs** | 10,465 | 11,719 | 11,712 | 11,708 |
| **RMSD** | 3.05 | 3.30 | 3.56 | 1.73 |
| **Alignment length** | 35.12 | 33.85 | 38.08 | 14.86 |
| **Normalized alignment length (%)** | 51.31 | 60.11 | 61.94 | 25.90 |
| **TM-score** | 0.32 | 0.28 | 0.31 | 0.23 |
| **Fraction of correctly aligned regions** | 41.70 | 41.34 | 41.30 | 40.50 |
| **Sequence similarity between structures** | 0.36 | 0.36 | 0.36 | 0.36 |
| **Fraction of comparisons for which a solution is not given for (%)** | 11.56 | 0 | 0 | 0.02 |
| **Time** **per alignment (s)** | 0.216 | 0.171 | 0.049 | 0.440 |

**Table S9.** Comparative metrics for MALISAM comparisons

| **MALISAM** | **Mican** | **SPalign-NS** | **TM-align** | **Multiprot** |
| --- | --- | --- | --- | --- |
| **Number of aligned pairs** | 80 | 108 | 107 | 75 |
| **RMSD** | 3.34 | 2.86 | 3.15 | **1.89** |
| **Alignment length** | 48.03 | 62.07 | **64.34** | 48.03 |
| **Normalized alignment length (%)** | **0.83** | 0.81 | **0.83** | 0.64 |
| **TM-score** | 0.47 | **0.53** | **0.53** | 0.26 |
| **Fraction of correctly aligned regions** | **60.19** | 58.98 | 56.72 | 46.25 |
| **Sequence similarity between structures** | 0.41 | 0.41 | 0.41 | 0.41 |
| **Fraction of comparisons for which a solution is not given for (%)** | 0 | 0 | 0 | 0 |
| **Time** **per alignment (s)** | 0.63 | 0.63 | **0.05** | 0.16 |

**Table S10.** Comparative metrics for MALISAM-ns comparisons

| **MALISAM-ns** | **Mican** | **SPalign-NS** | **TM-align** | **Multiprot** |
| --- | --- | --- | --- | --- |
| **Number of aligned pairs** | 74 | 63 | 63 | 71 |
| **RMSD** | 3.28 | 2.82 | 3.18 | **1.90** |
| **Alignment length** | 48.58 | 47.68 | **49.36** | 48.58 |
| **Normalized alignment length (%)** | **0.81** | 0.64 | 0.67 | 0.64 |
| **TM-score** | **0.52** | 0.42 | 0.42 | 0.28 |
| **Fraction of correctly aligned regions** | **68.44** | 55.86 | 55.50 | 49.36 |
| **Sequence similarity between structures** | 0.40 | 0.40 | 0.40 | 0.40 |
| **Fraction of comparisons for which a solution is not given for (%)** | 0 | 0 | 0 | 0 |
| **Time** **per alignment (s)** | 0.21 | 0.64 | **0.05** | 0.10 |

**Table 11.** Comparative metrics for MALIDUP comparisons

| **MALIDUP** | **Mican** | **SPalign-NS** | **TM-align** | **Multiprot** |
| --- | --- | --- | --- | --- |
| **Number of aligned pairs** | 189 | 228 | 231 | 213 |
| **RMSD** | 2.83 | 2.34 | 2.68 | **1.62** |
| **Alignment length** | 73.35 | 86.84 | **90.20** | 73.35 |
| **Normalized alignment length (%)** | **0.86** | 0.84 | **0.86** | 0.71 |
| **TM-score** | 0.60 | 0.64 | **0.65** | 0.36 |
| **Fraction of correctly aligned regions** | **81.48** | 77.22 | 76.55 | 70.16 |
| **Sequence similarity between structures** | 0.41 | 0.41 | 0.41 | 0.41 |
| **Fraction of comparisons for which a solution is not given for (%)** | 0 | 0 | 0 | 0 |
| **Time** **per alignment (s)** | 0.9 | 0.77 | 0.1 | 0.27 |

**Table S12.** Comparative metrics for MALIDUP-ns comparisons

| **MALIDUP-ns** | **Mican** | **SPalign-NS** | **TM-align** | **Multiprot** |
| --- | --- | --- | --- | --- |
| **Number of aligned pairs** | 219 | 179 | 178 | 218 |
| **RMSD** | 2.79 | 2.27 | 2.73 | **1.58** |
| **Alignment length** | 42.91 | 60.12 | **64.43** | 42.91 |
| **Normalized alignment length (%)** | **0.85** | 0.60 | 0.63 | 0.43 |
| **TM-score** | **0.58** | 0.46 | 0.47 | 0.36 |
| **Fraction of correctly aligned regions** | **79.53** | 74.29 | 70.85 | 70.70 |
| **Sequence similarity between structures** | 0.4 | 0.4 | 0.4 | 0.4 |
| **Fraction of comparisons for which a solution is not given for (%)** | 0 | 0 | 0 | 0 |
| **Time** **per alignment (s)** | 0.12 | 0.76 | **0.06** | 0.22 |

#### **3.4. TM-align and MultiProt comparison reveals TM-align aligns longer structures with comparable RMSD**

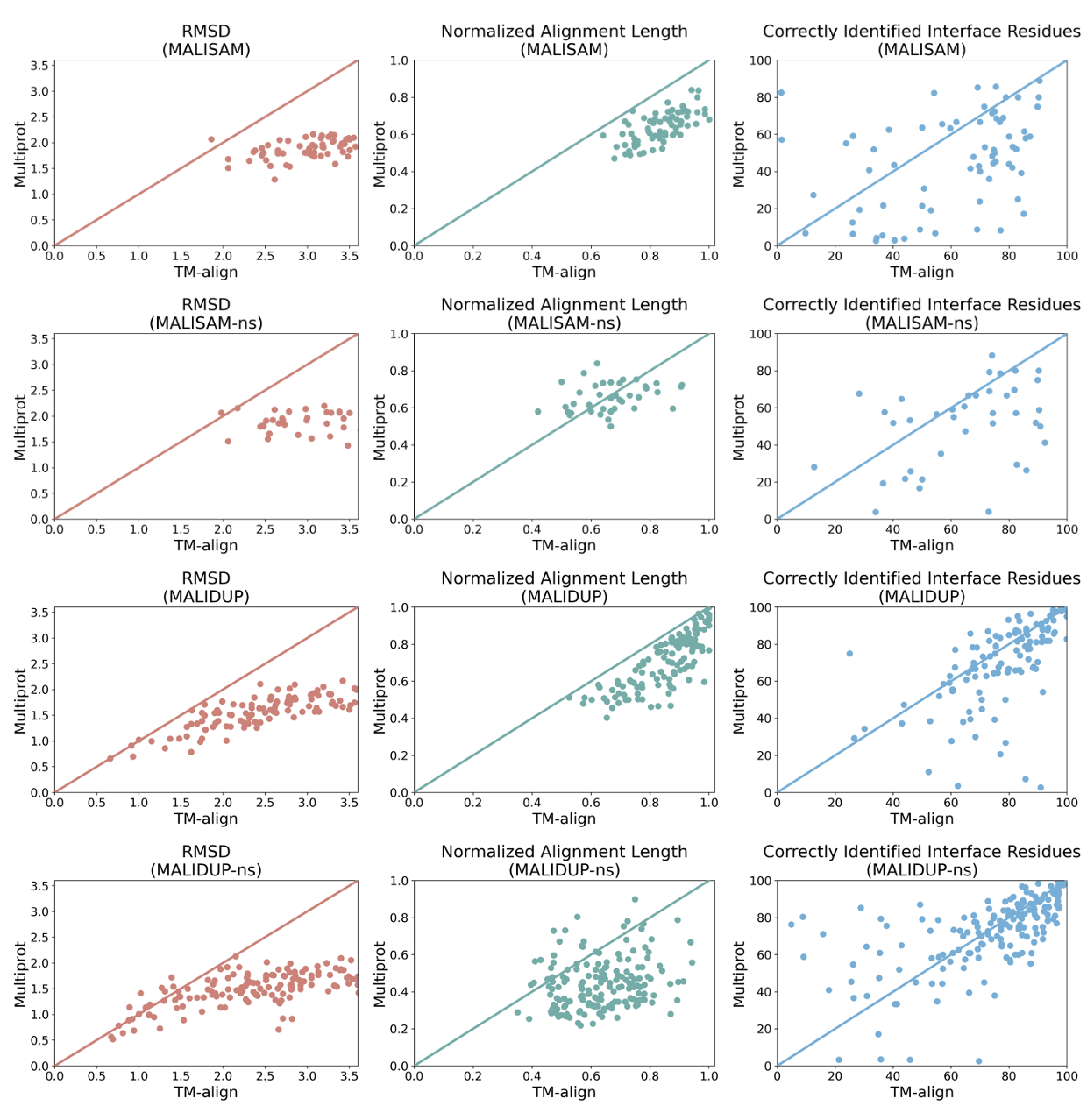

**Figure S4.** RMSD, normalized alignment length, and the fraction of correctly aligned residue comparisons for the subset pairs of benchmark sets for which results are given by both TM-align and MultiProt.
